# Quantifying invasive pest dynamics through inference of a two-node epidemic network model

**DOI:** 10.1101/2023.01.30.526176

**Authors:** Laura E Wadkin, Andrew Golightly, Julia Branson, Andrew Hoppit, Nick G Parker, Andrew W Baggaley

## Abstract

Invasive woodland pests are having a substantial ecological, economic and social impact, harming biodiversity and ecosystem services. Mathematical modelling informed by Bayesian inference can deepen our understanding of the fundamental behaviours of invasive pests and provide predictive tools for forecasting the future spread. A key invasive pest of concern in the UK is the oak processionary moth (OPM). OPM was established in the UK in 2006, is harmful to both oak trees and humans, and its infestation area is continually expanding. Here, we use a computational inference scheme to estimate the parameters for a two-node network epidemic model to describe the temporal dynamics of OPM in two geographically neighbouring parks (Bushy Park and Richmond Park, London). We show the applicability of such a network model to describing invasive pest dynamics and our results suggest that the infestation within Richmond Park has largely driven the infestation within Bushy Park.

## 1. Introduction

The spread of invasive pests, including non-native insects, within urban treescapes, woodlands and forests is having a profound environmental, economic and social impact [1–3]. In the UK, invasive species are estimated to have cost £5–13 billion since 1976 through damages and management costs, and impacts on ecosystem services [4]. Thus, the UK government has identified enhancing biosecurity as a key priority, through the control of existing pests and by building resilience against emerging concerns, harnessing computational modelling methods [5].

Statistical and mathematical computational models can be used to explore the fundamental behaviours of pest infestations and facilitate quantitative predictions of the future spread. Commonly employed models include dynamical systems models consisting of differential equations that describe pest population numbers and movements across landscapes, such as those for the grey squirrel in Wales [6]. Stochastic epidemic models [7] are more commonly used to describe disease dynamics within a population, but are transferable to the application of invasive pest spread [8]. In our previous work [9], we demonstrated that such an approach was indeed transferable to exploring invasive pest dynamics, and we build upon that framework here.

A key invasive pest of concern for treescapes within the UK and northern mainland Europe is the oak processionary moth (OPM), *Thaumetopoea processionea*. First established in the UK in 2006 through an accidental import, OPM is destructive to oak trees, causing defoliation which can leave infested trees vulnerable to other stressors. Additionally, OPM larvae have poisonous hairs which contain a urticating toxin harmful to both human and animal health [10,11]. Despite great efforts to contain the UK infestation to the originally affected area of south-east England [11], the extent of OPM continues to spread, with an expansion rate estimated at 1.7 km/year for 2006–2014, with an increase to 6 km/year from 2015 onwards [12]. There is evidence to suggest that the regions surrounding the current infestation area are particularly climatically suitable [13] and thus the prediction and control of the OPM population at its outer extent is especially crucial. Previous models for OPM have included species distribution models to predict the future infestation under climate change [13] and electric network theory models to predict high-risk regions [14].

In our previous work [9], we considered the temporal population of OPM in two London parks (Bushy Park and Richmond Park), applying a novel Bayesian inference scheme to estimate the parameters for a compartmental epidemic model with a time varying infestation rate. This showed that the infestation rate in both parks had remained relatively constant between 2013 and 2021, despite the control methods in place, resulting in the observed continual expansion of the infestation area [12].

In this paper, we build upon the work in [9], challenging the assumption that the infestations within the two parks are independent of each other, due to their neighbouring geographical location. Thus, here we consider a two-node compartmental epidemic model, using analogous computational inference methods (making use of a linear Gaussian approximation to the stochastic susceptible-infected-removed (SIR) model and a Markov chain Monte Carlo scheme) to estimate the infestation parameters, including a quantification of each park’s influence upon its neighbour. The data, model and inference scheme are detailed in Section 2, with the results presented in Section 3 and further discussion in Section 4. Our findings demonstrate the applicability of a two-node compartmental model to describe OPM spread and provide a framework applicable to other partially observed time series for infestations.

## 2. Materials and Methods

In this section, we present the observational OPM data (Section 2.1), detail the two-node SIR epidemic model (Section 2.2) and outline the statistical methods used to estimate the model parameters (Section 2.3 and 2.4).

### 2.1. The data

We consider the OPM infestation within two London parks: Bushy Park and Richmond Park. The data consists of the numbers and locations (eastings and northings) of removed OPM nests between 2013 and 2021. The data are collected and processed by The Royal Parks and shared with the Forestry Commission to inform the national OPM Control Programme. The University of Southampton (GeoData) provide analysis, support, and hold the data on behalf of the Forestry Commission.

A summary of OPM nest presence (and immediate removal) in both parks is shown in Figure 1(a) and (b). We consider the cumulative time series of previously infested trees (those recorded as having nests removed) as our observed data in the following sections, shown in Figure 1(c). This corresponds to the ‘Removed’ category prevalence in the compartmental SIR model presented in the next section, as in [9]. We refer to this as a partially observed dataset as we only have information about one of the categories in the compartmental SIR model. The neighbouring geographical location of the two parks, shown in Figure 1(d), motivates our choice to consider a two-node epidemic network model, outlined in the next section.

**Figure 1.**
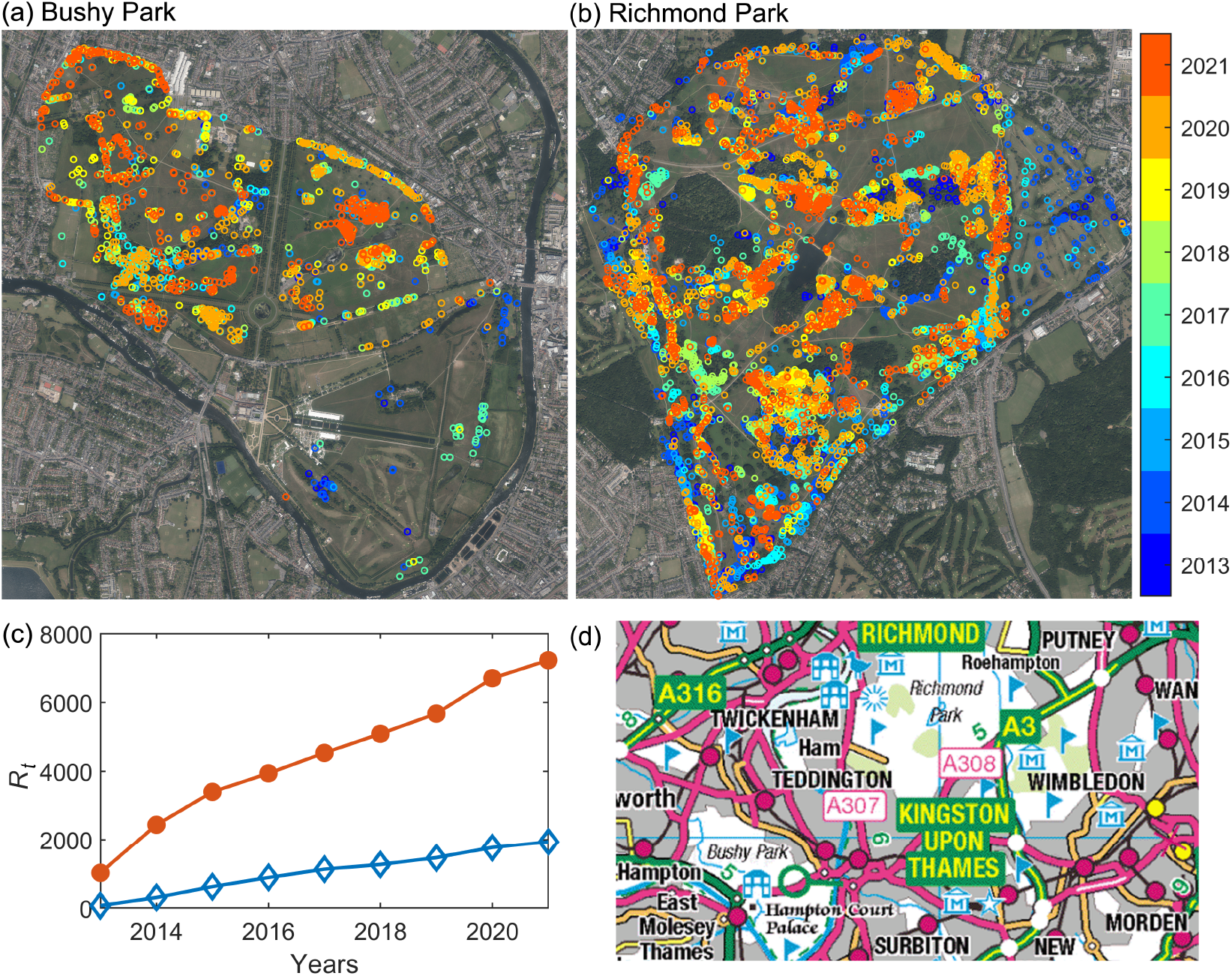
Satellite images (obtained from EDINA Digimap Aerial © Getmapping Plc [15]) of (a) Bushy Park and (b) Richmond Park with locations of nest removals between 2013 and 2021. (c) The ‘Removed’ prevalence time series, *R*_*t*_ (the cumulative numbers of tree locations where nest removal has taken place) is used as our observational data set. Bushy Park is shown in blue with open diamonds and Richmond park is shown in orange with filled circles. (d) A subsection of the Ordnance Survey map (OS Open Data [16]) showing the location of the two parks.

### 2.2. The two-node model

In [9] we applied a stochastic SIR epidemic model [17] to the spread of OPM in Bushy Park and Richmond Park between 2013 and 2021. Here we expand this model, noting that the two parks are close in geographical location, as shown in Figure 1(d), and thus the OPM dynamics within each park may not be independent of each other. We therefore consider a similar stochastic SIR model, but for two connected nodes (here representing each of the two parks) as illustrated in Figure 2.

**Figure 2.**
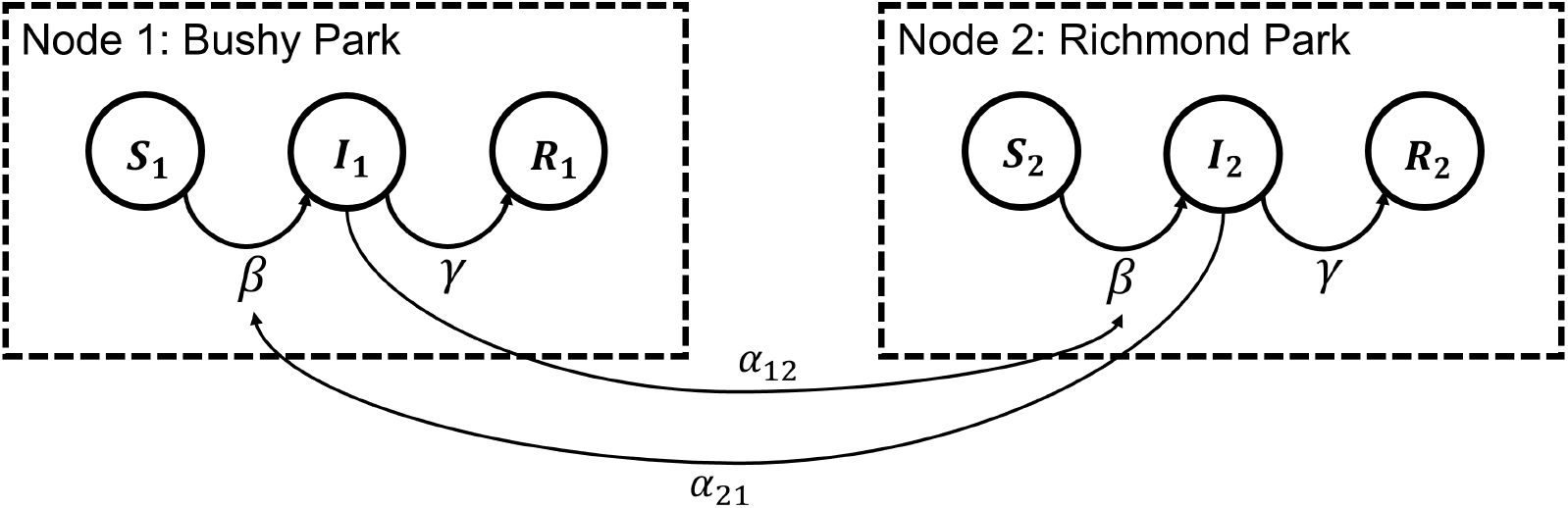
Illustration of the two-node epidemic network model. In each node the population of trees transition through the available states: Susceptible (*S*), Infested (*I*) and Removed (*R*). The transition between *I* and *R* is governed by the removal rate parameter, *γ*, in this case chosen to be identical within each node. For the transition between *S* and *I*, there is the standard infestation rate parameter, *β*, plus an additional infestation pressure resulting from the other node, with *α*_12_ corresponding to the infestation pressure from node 1 on node 2, and *α*_21_ the infestation pressure from node 2 on node 1. The resulting stochastic differential equation model for this scenario is given in Eqs (1)–(5).

Within each node, the fixed population of trees transition between the compartmental states: *S*, susceptible (not yet infested), *I*, infested (currently infested), and *R* removed (no longer infested or contributing to the infestation spread). The transition between the infested and removed state is governed by the removal rate parameter, *γ*. The transition between the *S* and *I* compartments is governed not only by the standard infestation rate parameter *β* (the rate at which contact of one infested tree with one susceptible tree will result in infestation, referred to as the ‘effective contact rate’), but also an additional infestation ‘pressure’ from the neighbouring node, described by the parameters *α*_*ij*_, where *α*_12_ is the pressure applied by node 1 on node 2, and *α*_21_ the pressure on node 2 by node 1. A similar model has been previously proposed to describe national surveillance counts from the 2013–2015 West Africa Ebola outbreak [18].

We assume that the effective contact rate *β*, and the removal rate *γ* are the same in both parks as parameters inherent to the OPM population under similar conditions. The effective contact rate *β* can be expressed as *β* = *κτ*/*N*, where *κ* is the number of contacts (opportunities for transmission) and *τ* is the transmissibility of the disease (here, the pest). Since *τ* is inherent to the system, the assumption of equal effective contact rates in each node results in *κ*_1_/*N*_1_ = *κ*_2_/*N*_2_ and thus a number of contacts *κ* that is proportional to the population size within each node.

The dynamics of all compartment states in the two-node model above are most naturally described by a Markov jump process (MJP) whereby state numbers are described via a continuous-time Markov process with a discrete state-space, reflecting the fact that states change abruptly and discretely in time [7]. As noted in [19] this can be computationally prohibitive for models in which typical population sizes are more than a few hundred. We therefore eschew the MJP representation in favour of a tractable continuous approximation via a stochastic differential equation (SDE). We describe the SDE below before considering a further tractable approximation known in the stochastic kinetics literature as the linear noise approximation (LNA). We refer the reader to [20] for further details on the SDE and LNA approximation of an MJP.

The corresponding stochastic differential equation model considers the latent process *X*_*t*_ = (*S*_1,*t*_, *I*_1,*t*_, *S*_2,*t*_, *I*_2,*t*_)^′^, where *S*_*i,t*_ and *I*_*i,t*_ denote the number of trees in each of the compartments *S* and *I* in node *i* at time *t* ≥ 0. The complete SDE model can be describedby

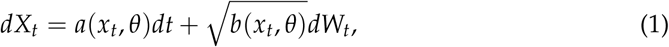

where *x*_*t*_ = (*s*_1,*t*_, *i*_1,*t*_, *s*_2,*t*_, *i*_2,*t*_) is the state of the system at time *t, θ* = (*β, γ, α*_12_, *α*_21_)^′^ is the vector of parameter values and *dW*_*t*_ = (*W*_1,*t*_, *W*_2,*t*_, *W*_3,*t*_, *W*_4,*t*_)^′^ denotes uncorrelated standard Brownian motion processes on each of the compartmental states. The SDE drift function *a*(*x*_*t*_, *θ*) and diffusion coefficient *b*(*x*_*t*_, *θ*) are given by

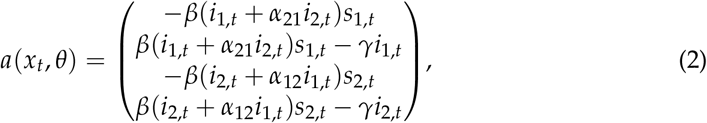

and

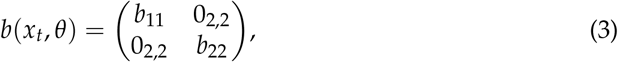

where

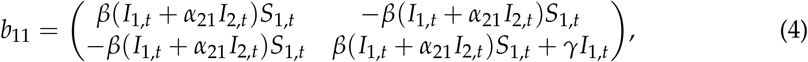

and

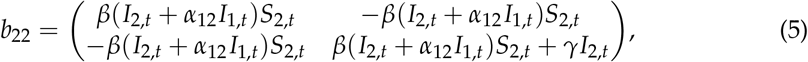

and 0_2,2_ is the 2 × 2 zero matrix. Since the SDE specified by (1)–(5) cannot be solved analytically, we replace the intractable analytic solution with a tractable Gaussian process approximation: the LNA, described in the next section.

### 2.3. Linear noise approximation

The LNA, provides a tractable approximation to the SDE given in (1)–(5). We used the LNA in the same manner for a stochastic SIR model with a time varying infestation rate, also applied to OPM, in [9]. Formal details of the LNA can be found in [21–23]; below we outline the derivation.

Consider a partition of *X*_*t*_ as

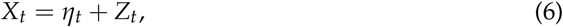

where {*η*_*t*_, ≥ *t* 0} is a deterministic process satisfying the ordinary differential equation (ODE)*dt*

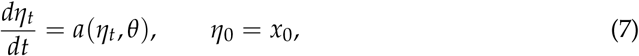

and {*Z*_*t*_, *t* ≥ 0} is a residual stochastic process. The residual process *Z*_*t*_ satisfies

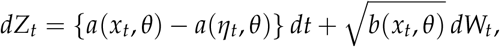

which will typically be intractable. The assumption that || *X*_*t*_− *η*_*t*_|| is “small” motivates a Taylor series expansion of *a*(*x*_*t*_, *θ*) and *b*(*x*_*t*_, *θ*) about *η*_*t*_, with retention of the first two terms in the expansion of *a* and the first term in the expansion of *b*. This gives an approximate residual process 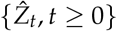 satisfying

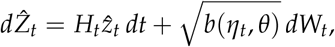

where *H*_*t*_ is the Jacobian matrix with (*i,j*)th element

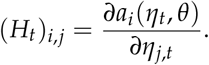

For the SIR model in (1)–(5) we therefore have

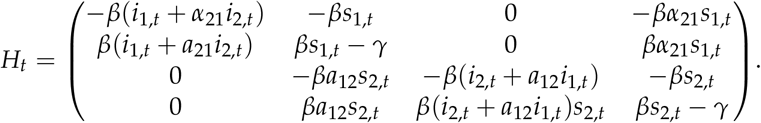

Given an initial condition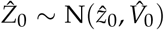, it can be shown that 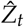 is a Gaussian random variable [24]. Consequently, the partition in (6) with *Z*_*t*_ replaced by 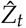, and the initial conditions *η*_0_ = *x*_0_ and 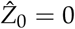 give

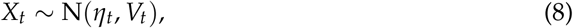

where *η*_*t*_ satisfies (7) and *V*_*t*_ satisfies

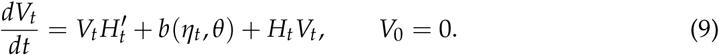

Further details on the derivation of (9) are given in [9]. Hence, the linear noise approximation is characterised by the Gaussian distribution in (8), with mean and variance found by solving the ODE system given by (7) and (9), which can be solved numerically.

### 2.4. Bayesian inference

We consider the case in which not all components of the stochastic epidemic model are observed and that the data points are subject to measurement error, as in [9]. Observations (on a regular grid) *y*_*t*_, *t* = 0, 1, … *n* are assumed conditionally independent (given the latent process *X*_*t*_) with conditional probability distribution obtained via the observation equation,

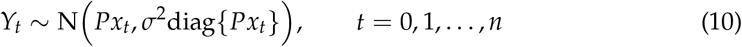

where

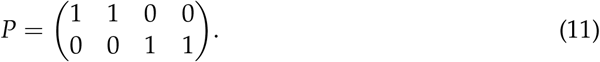

This choice of *P* is due to the data consisting of observations on the removed states in each node, *R*_1,*t*_ and *R*_2,*t*_, which for known population sizes *N*_1_ and *N*_2_, is equivalent to observing the sums *S*_1,*t*_ + *I*_1,*t*_ and *S*_2,*t*_ + *I*_2,*t*_. Our choice of observation model is motivated by a Gaussian approximation of two independent Poisson random variables with rates given by the components of *Px*_*t*_. Moreover, the assumption of a Gaussian observation model admits a tractable observed data likelihood function, when combined with the LNA (see Section 2.3 and [24,25]) as a model for the latent epidemic process *X*_*t*_. Details on a method for the efficient evaluation of this likelihood function can be found in [9].

Given data *y* = (*y*_0_, *y*_1_, …, *y*_*n*_) and upon ascribing a prior density *π*(*θ*) to the components of *θ* = (*β, γ, α*_12_, *α*_21_, *σ*)^′^, Bayesian inference proceeds via the joint posterior for the static parameters *θ* and unobserved dynamic process *x* = (*x*_0_, *x*_1_, …, *x*_*n*_). We have that

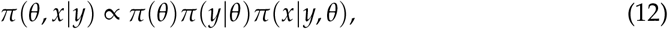

where *π*(*y*| *θ*) is the observed data likelihood and *π*(*x y*, | *θ*) is the conditional posterior density of the latent dynamic process. We use a Markov chain Monte Carlo scheme for generating (dependent) samples from (12) due to the intractable joint posterior. Briefly,| this comprises two steps: i) the generation of samples *θ*^(1)^, …, *θ*^(*M*)^ from the marginal parameter posterior *π*(*θ*| *y*) ∝ *π*(*θ*)*π*(*y*| *θ*) and ii) the generation of samples *x*_(1)_, …, *x*_(*M*)_ by drawing from the conditional posterior *π*(*x y, θ*^(*i*)^), *i* = 1, …, *M*.

The parameters required as input for the inference scheme are given in Table 1. We take estimates of the numbers of trees in each park and the infestation intitial conditions as in [9], with *N*_1_ = 5000 (Bushy) and *N*_2_ = 40000 (Richmond) and initial ODE conditions *x*_0_ = (*S*_1,0_, *I*_1,0_, *S*_2,0_, *I*_2,0_) = (*N*_1_ − *I*_1,0_ − *R*_1,0_, *I*_1,0_, *N*_2_ − *I*_2,0_ − *R*_2,0_, *I*_2,0_).

**Table 1.** The input parameters used in the inference scheme for the two-node model (Section 2.2– 2.4).

| Input parameters |  |
| --- | --- |
| Total population | $N_1 = 5000, N_2 = 40000$ |
| Initial infested | $I_{1,0} = 240, I_{2,0} = 1400$ |
| Initial model parameters | $\theta_0 = (\beta_0, \gamma_0, \alpha_{12_0}, \alpha_{21_0}, \sigma_0) = (0.00001, 0.8, 0.3, 2.5, 1)$ |
| Prior distributions | $\log \theta_i \sim N(0, 1^2), \quad i = 1, \dots, 4$ |
| Tuning parameter | $\Sigma = \begin{pmatrix} 0.009 & 0.003 & -0.059 & -0.009 & 0.002 \\ 0.003 & 0.006 & 0.011 & 0.001 & 0.001 \\ -0.059 & 0.011 & 0.712 & 0.072 & -0.006 \\ -0.009 & 0.001 & 0.072 & 0.013 & -0.002 \\ 0.002 & 0.001 & -0.006 & -0.002 & 0.052 \end{pmatrix}$ |
| Initial ODE mean | $x_0 = \eta_0 = (N_1 - I_{1,0} - R_{1,0}, I_{1,0}, N_2 - I_{2,0} - R_{2,0}, I_{2,0})$ |
| Initial ODE variance | $V_0 = 0_{4,4}$ |
| Observation matrix | $P = \begin{pmatrix} 1 & 1 & 0 & 0 \\ 0 & 0 & 1 & 1 \end{pmatrix}$ |

## Results

We take the data, detailed in Section 2.1 and pictured in Figure 1, for the cumulative number of trees with removed OPM nests in Bushy Park and Richmond Park. We (arbitrarily) denote Bushy Park as node 1, and Richmond Park as node 2, with the observed data corresponding to the removal prevalence time series *R*_1,*t*_ and *R*_2,*t*_, in the two-node model described in Section 2.2. Through the inference techniques outlined in Section 2.3 and Section 2.4, we infer the parameters for the two-node stochastic epidemic model: the infestation rate *β*, and removal rate *γ*, common to both nodes, along with additional parameters representing the infestation ‘pressure’ resulting from the neighbouring node, *α*_12_ and *α*_21_, and an observation error *σ*.

The results from the inference scheme are shown in Figure 3 with within-sample median posterior series for *S*_*i,t*_, *I*_*i,t*_ and *R*_*i,t*_, and posterior densities of the inferred parameters. The average parameter results are shown in full in Table 2. The median infestation rate is *β* ≈ 1.8× 10^−5^, the median removal rate is *γ* ≈ 0.8, and the median observation error (see (10)) is *σ* ≈1.3. The median posterior estimates of the parameters connecting the two nodes are *α*_12_ ≈ 0.3 (Bushy to Richmond) and *α*_21_ 3 (Richmond to Bushy).

**Table 2.**
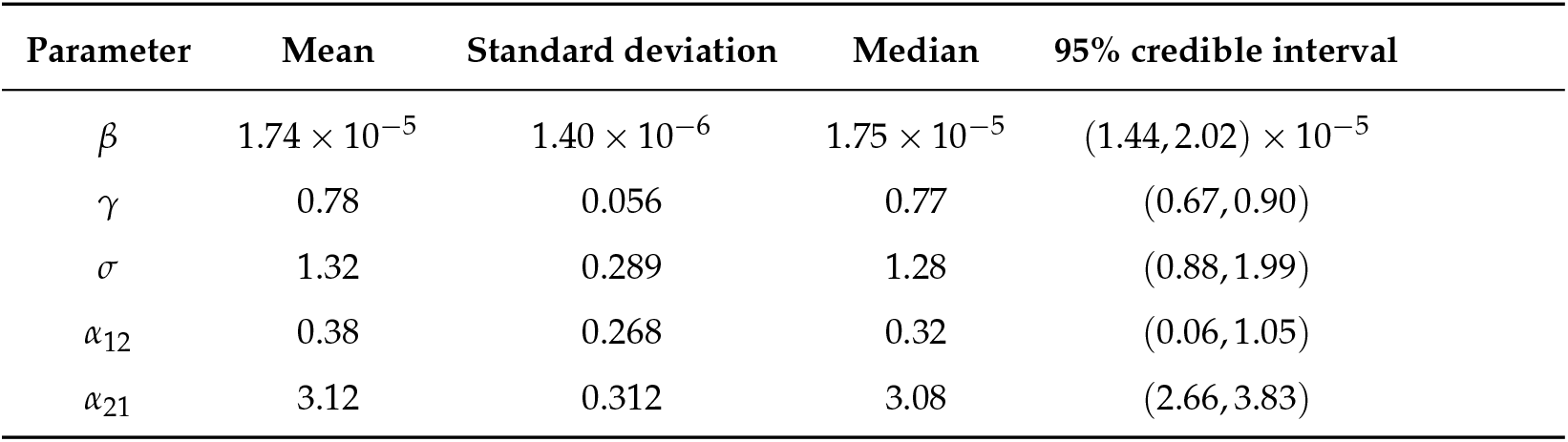
The posterior means, standard deviations, medians and 95% credible intervals for the inferred model parameters.

| Parameter | Mean | Standard deviation | Median | 95% credible interval |
| --- | --- | --- | --- | --- |
| $\beta$ | $1.74 \times 10^{-5}$ | $1.40 \times 10^{-6}$ | $1.75 \times 10^{-5}$ | $(1.44, 2.02) \times 10^{-5}$ |
| $\gamma$ | 0.78 | 0.056 | 0.77 | (0.67, 0.90) |
| $\sigma$ | 1.32 | 0.289 | 1.28 | (0.88, 1.99) |
| $\alpha_{12}$ | 0.38 | 0.268 | 0.32 | (0.06, 1.05) |
| $\alpha_{21}$ | 3.12 | 0.312 | 3.08 | (2.66, 3.83) |

**Figure 3.**
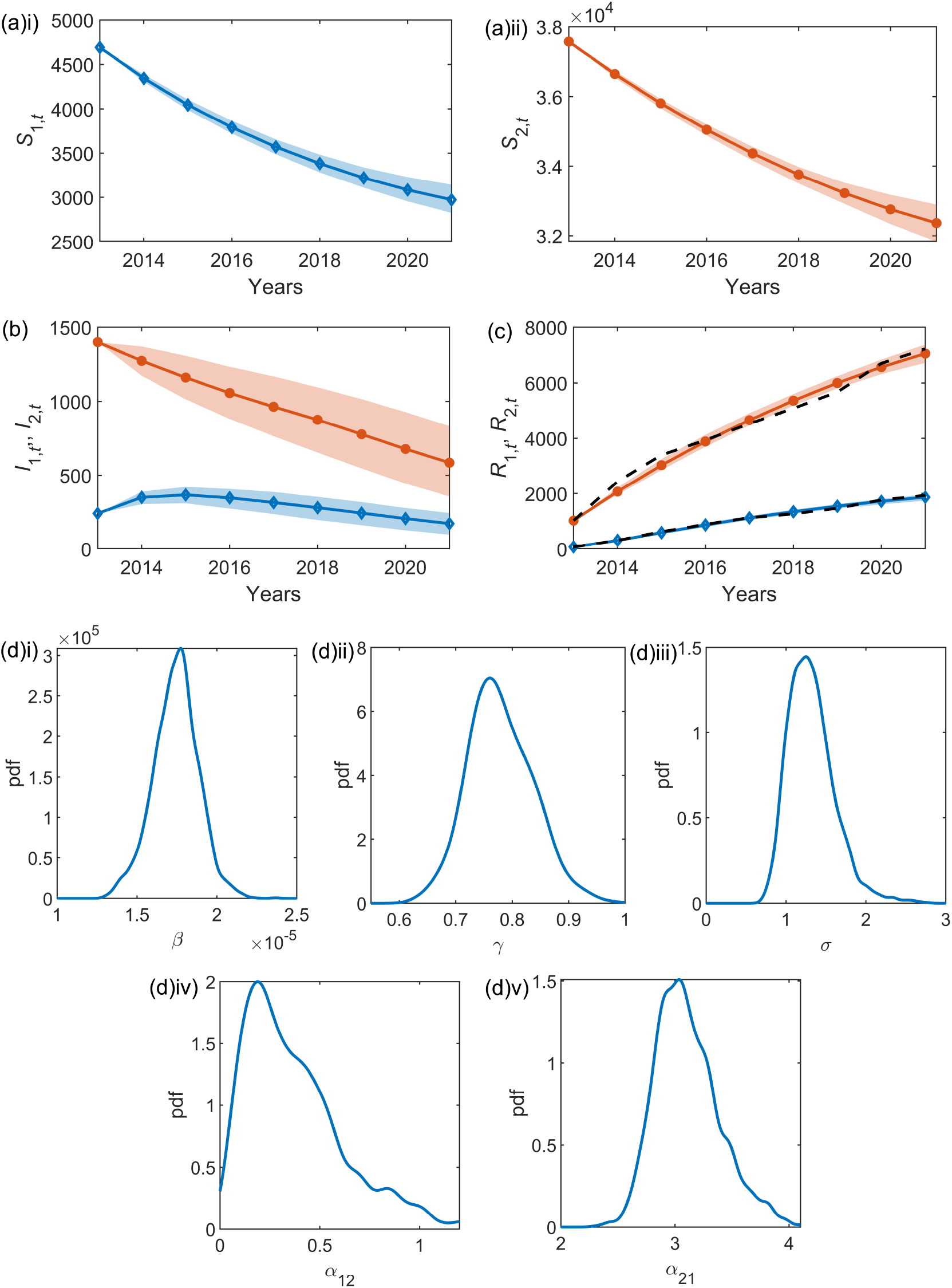
Inference results for the two-node network model applied to Bushy and Richmond Park. The within-sample posteriors for (a) the susceptible tree time series for i) Bushy Park, *S*_1,*t*_, and ii) Richmond Park, *S*_2,*t*_, (b) the infested tree time series, *I*_1,*t*_ and *I*_2,*t*_, and (c) the removed tree time series, *R*_1,*t*_ and *R*_2,*t*_. The median of the within-sample posteriors are shown for Bushy Park with blue diamonds and Richmond Park with orange circles. In all cases, the shaded error bars represent the 95% credible interval. In (c) the observed time series for *R*_1,*t*_ and *R*_2,*t*_ are shown as the black dashed lines. In (d) the posterior parameter distributions are shown.

We can consider each of the infestation components (see (2)): the standard intra-park component *βI*_*i*_*S*_*i*_, and the connecting inter-park component *βα*_*ij*_ *I*_*i*_*S*_*j*_. Probability densities for both the intra- and inter-park infestation components using the median posterior estimates of *S*_*i,t*_ and *I*_*i,t*_, and the full posterior distributions of *β* and *α*_*ij*_, are shown in Figure 4. Similarly, the expected number of new infestations from each of the two components, averaged over 50 forward simulations with the median parameters estimated through the inference scheme (Table 2), are shown in Figure 5. Both Figure 4 and Figure 5 illustrate that in Bushy Park the inter-park dynamics (Richmond-Bushy) are significantly greater than the intra-park (Bushy-Bushy), whereas in Richmond Park the intra-park dynamics (Richmond-Richmond) are more significant than the inter-park (Bushy-Richmond). This suggests that the infestation in Bushy Park has been largely driven by the infestation in Richmond Park to a greater extent than vice versa.

**Figure 4.**
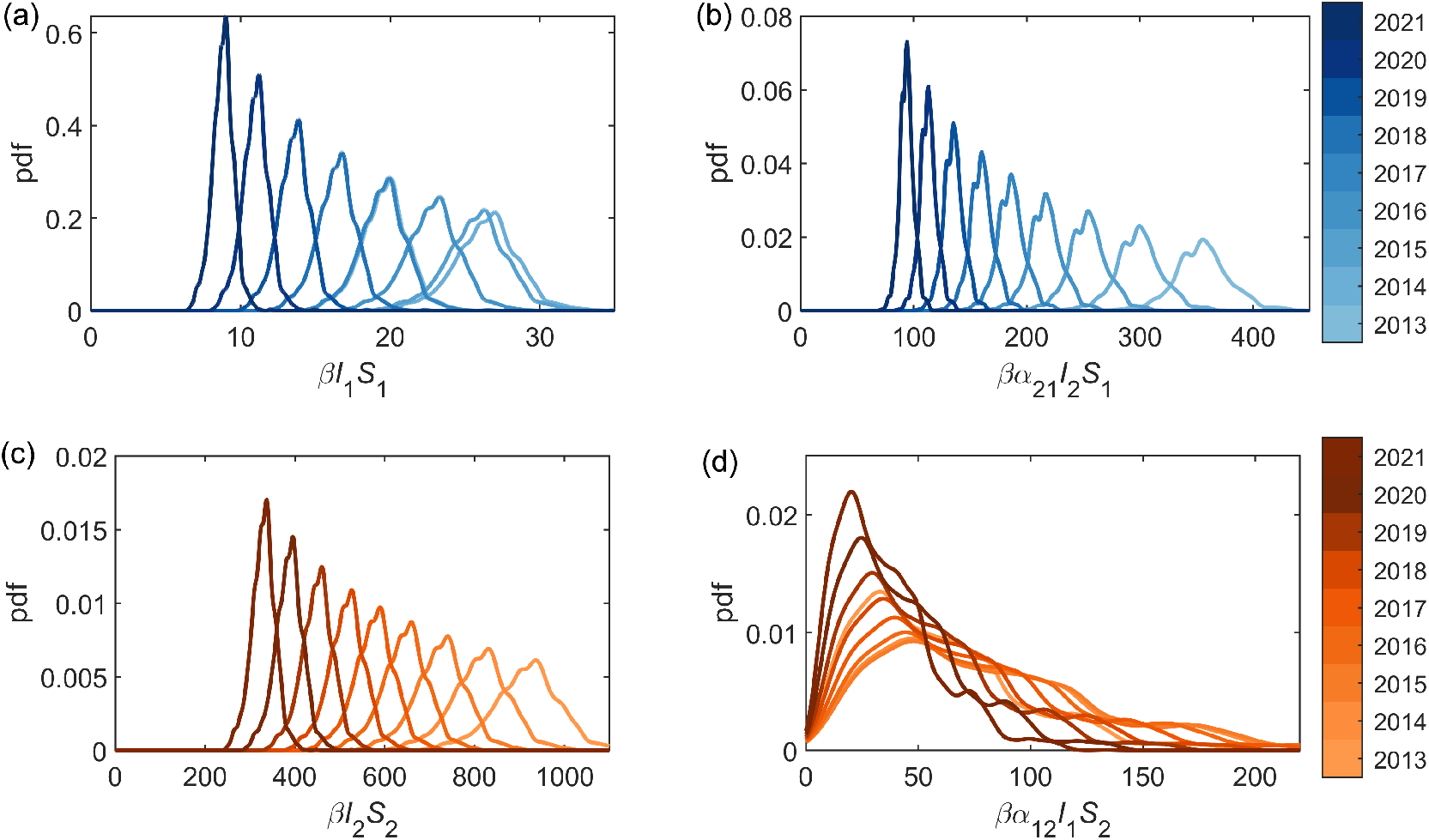
The estimated intra-park (*βI*_*i*_ *S*_*i*_) and inter-park (*βα*_*ij*_ *I*_*i*_ *S*_*j*_) infestation components (see (2)) using the median posterior estimations for *I* and *S* in each year for (a) and (b) Bushy Park, and (c) and (d) Richmond Park.

**Figure 5.**
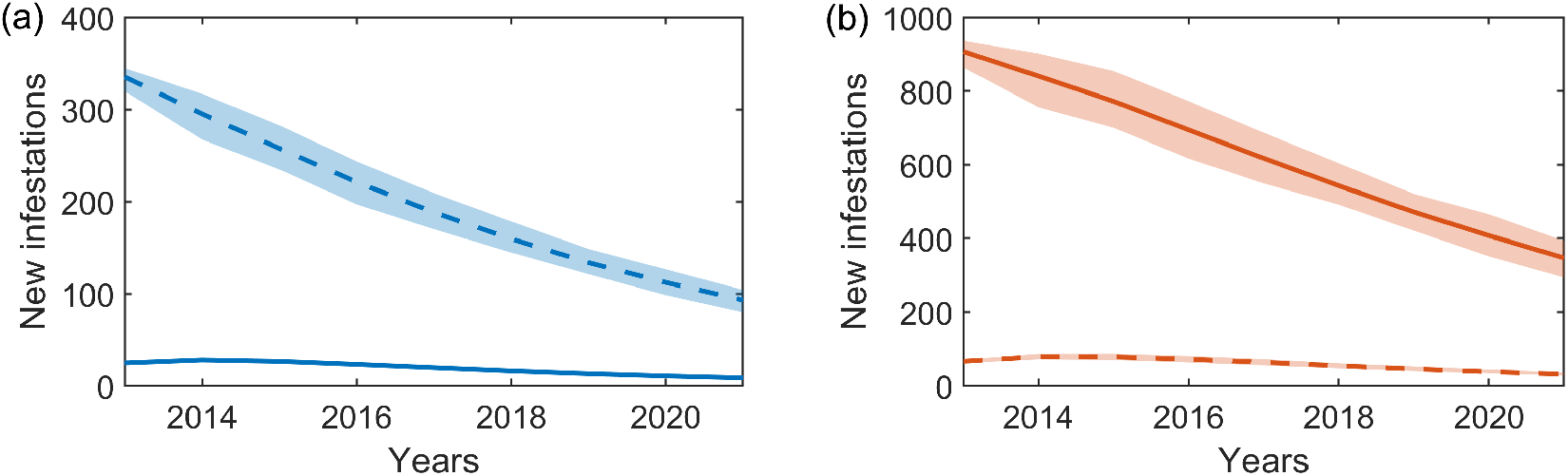
An example scenario for possible new infestations occurring in (a) Bushy and (b) Richmond Park, averaged over 50 simulations of the stochastic two-node epidemic model with the median parameters estimated through the inference scheme (shown in Table 2). In both panels, the solid lines show the new infestations resulting from within the park (intra-park), with dashed lines showing new infestations resulting from the neighbouring park (inter-park). Error bars show the 95% credible intervals over the 50 simulations.

Predictions of the spread of OPM as required to inform management strategies. We can use the inferred model parameters to simulate the infestation forwards in time. The simulated removal prevalence time series resulting from the stochastic two-node epidemic model with the median inferred parameter estimates is shown in Figure 6 for the years 2013 to 2025. Here we see the overall alignment with the observational data, with a deviation between 2017 and 2020 in which the infestation numbers were lower than this model predicts, and the future forecasting if the infestation were to continue with these characteristic parameters.

**Figure 6.**
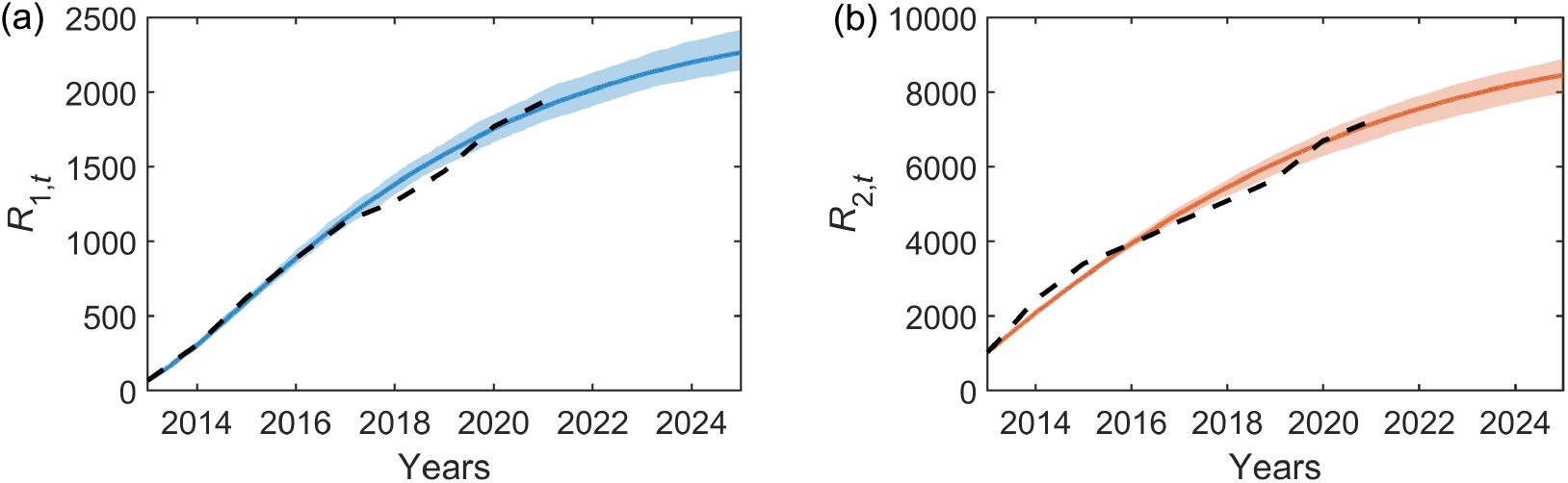
Forward simulations for Removal prevalence from the stochastic two-node epidemic model with the median parameters estimated through the inference scheme (shown in Table 2).

A possible contributing factor to the dynamics, not considered explicitly in this epidemic model, is the density of OPM nests within each park, i.e., the numbers of nests per infested tree, shown in Figure 7. Richmond park has a slightly higher infestation density, with a median of three nests per infested tree, compared to the median of two nests per tree in Bushy Park, and an upper quartile of six nests per tree, compared to the upper quartile of four in Bushy Park. This increased nest density could contribute to the resulting infestation pressure from the Richmond Park infestation to the Bushy Park infestation.

**Figure 7.**
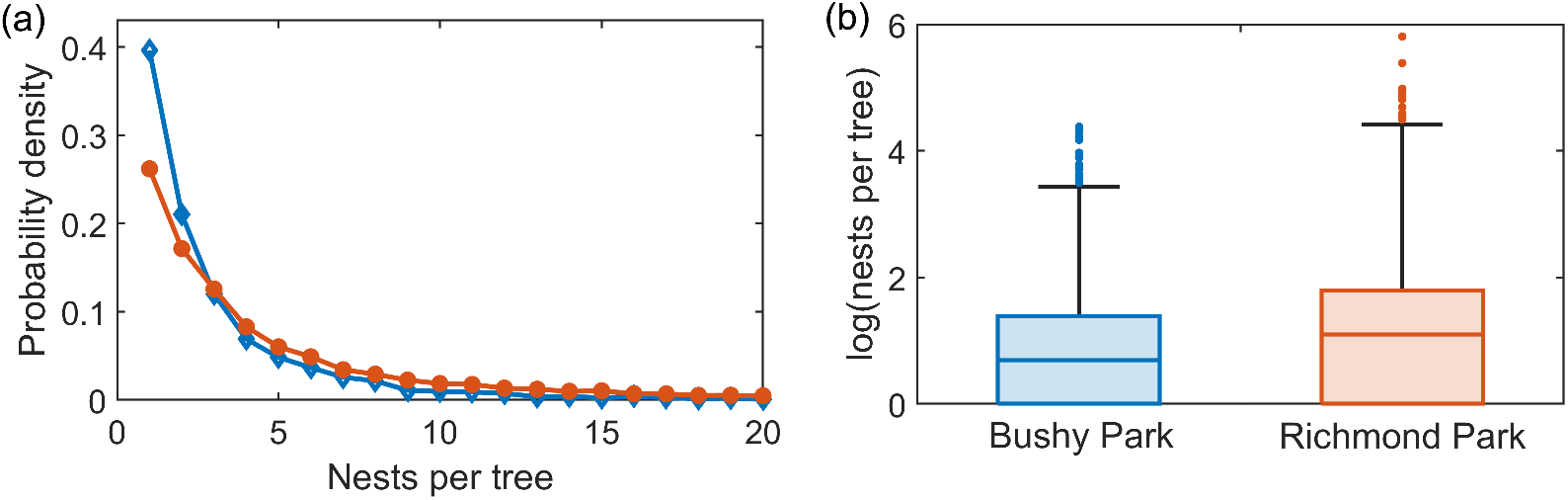
OPM nest density. (a) Probability density of the number of recorded removed nests per tree in Bushy Park (blue, open diamonds) and Richmond Park (orange, filled circles). For visualisation purposes this shows nest numbers up to 20 per tree only. (b) Box plots of the number of logged nests per tree on a logarithmic scale, showing all recorded data.

## 4. Discussion

It is crucial to deepen our understanding of the dynamics of invasive pests to develop predictive modelling tools and maximise the impact of control strategies. Here we have shown the applicability of two-node compartmental epidemic models, using case study data of the OPM infestations within the neighbouring Bushy Park and Richmond Park in London.

In our previous work [9] exploring the UK OPM infestation, we considered the two parks as independent contained areas and estimated the parameters for a compartmental model with a time-varying infestation rate, showing the infestation rate had remained stable over time. Here we challenge the assumption that the infestations within the two parks are independent due to their geographical proximity (with the closest park boundaries separated by approximately 2 km).

Instead, we assume that the infestation contact rate within each park, *β*, (assumed to be a constant based on the results from [9]) and the removal rate, *γ*, are inherent properties of the OPM species under similar conditions, resulting in identical *β* and *γ* in both nodes (parks). Since *β* can be expressed as *κτ*/*N*, with *κ* the number of contacts and *τ* a transmissibility parameter inherent to the species, we have *κ*_1_/*N*_1_ = *κ*_2_/*N*_2_, and thus that the number of contacts scales with total population number. In the case of OPM this corresponds to an increasing number of ‘contacts’ between trees due to the underlying movement of the OPM population, an assumption that would hold whilst the typical movement distances of OPM are of a similar (or greater) length scale to the tree population area, i.e., a greater number of trees (increased *N*) means a greater number of opportunities for contacts (increased *κ*) providing the moths can travel over the whole area. Despite the joint infestation parameters *β* and *γ*, the infestation dynamics can still differ in each park due to different numbers of susceptible trees and the introduction of two additional parameters connecting the infestations within the two nodes, *α*_12_ and *α*_21_.

Similar epidemic models have previously been used to describe the spread of infectious disease within human populations, e.g., for ebola in [18]. In these cases the parameters connecting the nodes represent the rates of the movement of individuals between nodes, e.g., the movement of infected people between geographical locations. In the case of our OPM model, the individuals within each of the SIR compartments are trees, which are spatially fixed, and thus the *α*_*ij*_ instead represent a proxy measure of the movement of the underlying OPM population between the two nodes. The movement patterns of OPM are not fully characterised, but can occur through three possible routes: short-distance larvae movement, flight of adult moths, and accidental human-mediated dispersal [12,26]. Considering the locations of the two parks considered here, the latter two dispersal mechanisms are possibilities for facilitating moth movement between nodes.

We find that the connecting parameter representing the infestation pressure on the trees in Bushy Park resulting from the infestation in Richmond park is much stronger than vice versa. The infestation in Richmond Park is relatively unaffected by the infestation in Bushy Park. One reason for the observed Richmond-led infestation dynamics could be the higher underlying nest density within the park, resulting in a greater population density of OPM per susceptible tree, a larger contact number, *κ*, and more opportunities for longer-range movement between the two parks. The relationship between OPM nest density, tree height, infestation percentages, and bacterial control treatment is explored in [27].

We note that there are control measures taking place in many OPM infested areas [28]. In Bushy and Richmond Parks, control measures include the yearly nest removal (leading to the data used here) and limited spraying with a biological insecticide which has been shown to reduce nest density [27]. Thus, all inferred parameters represent the infestation dynamics under these conditions, rather than inherent parameters of uncontrolled pest spread. Although not ideal for learning about the fundamental properties of the species, this is necessary as emergent invasive pests require an immediate control response.

Future work could explore expanding the epidemic network to a greater number of areas (nodes), forming more connections across the wider OPM infested area of south-east England. If a similar approach to network building was taken to that described in [14], the results from the statistical compartmental epidemic model could be compared with the electric network theory model [14]. However, expanding the network represents a computational challenge, with increasing numbers of parameters to infer.

The results from this work can inform the development of future computational models for the spread of OPM, and provide a statistical framework for applying to other emerging pest concerns with similar partially-observed temporal data sets.

## Author Contributions

Conceptualization, L.W., A.G., N.G.P. and A.B.; methodology, L.W., A.G., N.G.P. and A.B.; software, L.W. and A.G.; validation, L.W. and A.G.; formal analysis, L.W. and A.G.; writing—original draft preparation, L.W.; writing—review and editing, L.W., A.G., J.B., A.H., N.G.P. and A.B.; visualization, L.W.; funding acquisition, L.W., A.G., N.G.P. and A.B.. All authors have read and agreed to the published version of the manuscript.

## Data Availability Statement

All statistical modelling software with the Bushy and Richmond Park time series are available on the Newcastle University data repository at XXX (to be added upon publication). The full OPM data used in this paper were collected, processed, and kindly shared by The Royal Parks charity. This data can be provided from The Royal Parks upon reasonable request.

## Acknowledgments

We thank Gillian Jonusas from The Royal Parks for sharing the OPM data for Bushy and Richmond parks.

## Abbreviations

The following abbreviations are used in this manuscript:

LNA: Linear noise approximation MJP Markov jump process
ODE: Ordinary differential equation
OPM: Oak processionary moth
SDE: Stochastic differential equation
SIR: Susceptible, infected, removed

